# Slow trajectories generate divergent cell fates following antibiotic stress

**DOI:** 10.1101/2025.09.25.678360

**Authors:** Ethan Levien, João Pedro Teuber Carvalho, Linh Huynh, Daniel Schultz

## Abstract

Single-cell microfluidic experiments have shown that upon abrupt exposures to antibiotics, genetically homogeneous microbial populations undergo divergent cell fates. The mechanism underlying this divergence is not clear and in particular, the emergence of a range of distinct slow-growing phenotypes cannot be explained by models relying on bistability alone. Here, we propose a model for gene expression and growth dynamics during antibiotic exposures, which is informed by well-known scaling relations connecting proteome allocation and cell growth. In our model, resources available for transcription and translation of resistance genes act like generalized momenta and their initial variation is predictive of cell fate. Our model reproduces key experimental observations, including the prediction of specific phenotypes and a critical threshold in initial resource allocation that predicts cell survival. These results offer an alternative mechanism for the emergence of phenotypic diversity where slow cell growth effectively stabilizes cellular states along the trajectory that are far from the final stable states predicted by fixed points.

## Introduction

The question of how phenotypically diverse behaviors emerge within genetically identical cells has been at the center of microbiology for nearly 100 years, starting with the observation that subsets of microbial populations can adopt different phenotypes. In particular, persister *Staphylococcus aureus* cells that could tolerate penicillin treatments were identified within an otherwise susceptible population shortly after the advent of antibiotics [6, 31]. In this example, population heterogeneity is not the result of mutations and is instead attributed to the existence of multiple stable states within the underlying regulatory network [1, 4, 5, 13, 15, 28, 40, 48]. However, recent studies have observed that microbial populations under stress often exhibit phenotypic distributions which do not seem to result from a few well-defined stable states [17, 36, 41].

In addition to transcriptional regulation, antibiotic resistance phenotypes have also been linked to variation in cell metabolism [3, 7, 18, 19, 22, 24, 30, 35, 37]. In the last decade, several studies have established phenomenological relationships between drug action, cell growth, cellular resource allocation, and expresion of resistance genes [16, 23, 44, 46]. Antibiotic effects on cellular metabolism often slow cell growth and affect the allocation of resources towards the expression of several cellular responses, resulting in markedly different phenotypes (Fig. 1A) [45, 47]. Furthermore, slow growth also prevents cells from quickly reaching stable phenotypes, increasing the time spent in transient phenotypic states [14]. Therefore, metabolically active slow-growing cells can effectively stabilize alternative phenotypes far from equilibrium.

**FIG. 1.**
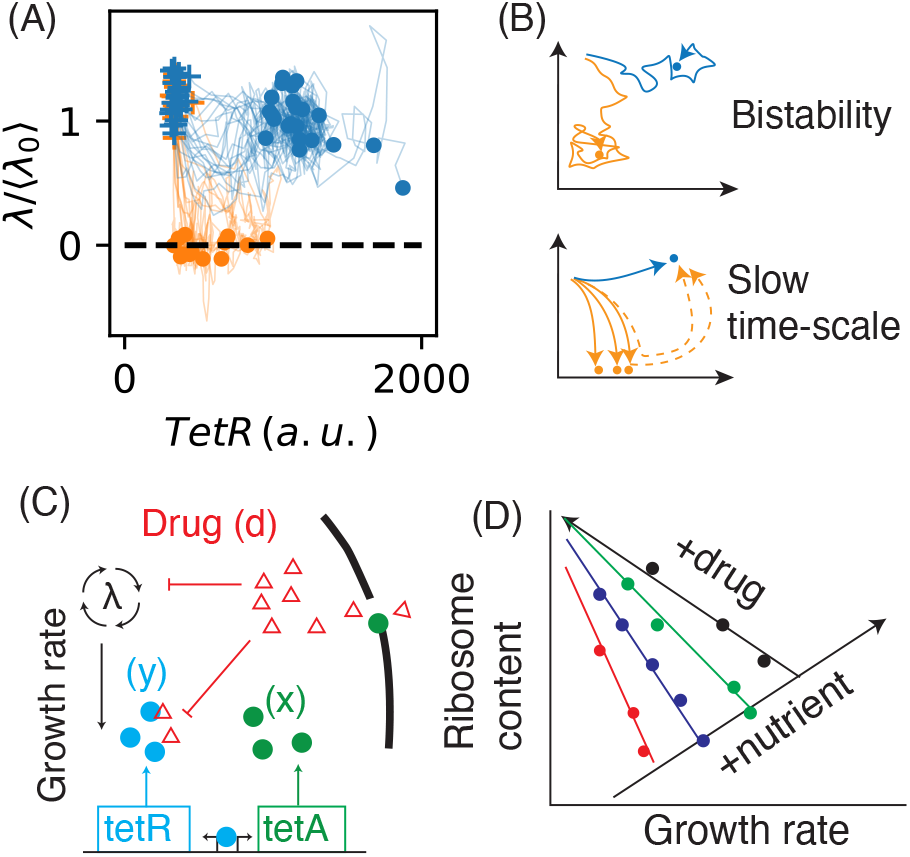
(A) Trajectories depicting expression of TetR and growth rates *λ* following and abrupt exposure to tetracycline, obtained from single-cell microfluidic experiments [42, 49]. Blue: recovered cells; Orange: Arrested or slow growing cells. Crosses indicate each growth rate prior to drug exposure and dots indicate time-averages after 3 generations under drug conditions. (B) Two potential mechanism for generating heterogeneous cell-fates. (top) Bistability, where cell-fates result from multiple fixed points in the gene expression dynamics. (bottom) Cell-fates result from a slow time-scale in the dynamics for certain initial conditions. (C) Diagram of the TetA-TetR system, showing the main biochemical interactions. (D) Linear scaling relations between ribosome content and cell growth rate, according to the theorem of the proteome partition [44].

Here, we present a model that illustrates how diverse cell fates can emerge through a mechanism that does not involve multistability. In our model, cells that do not express resistance fast enough become trapped along a slow trajectory toward the stable state (Fig. 1B). We base our model on a microfluidic experiment in which single *Escherichia coli* cells were suddenly exposed to the ribosome-inhibiting antibiotic tetracycline, initiating a response mediated by the expression of the TetA efflux pump, which exports tetracycline out of the cell (Fig. 1C) [11, 21, 29, 42]. In the absence of the antibiotic, the concentration of TetA is strongly repressed by the self-repressing regulator TetR [8, 20]. Upon exposure, tetracycline binds and inactivates TetR, releasing the expression of both *tet* genes [9, 38, 39].

This microfluidic experiment used fluorescent reporters to track cell growth and the expression of TetA, TetR, and a constitutive protein in single cells during exposures to tetracycline. Such study observed that drug exposures result in the coexistence of growing cells expressing TetA (recovered) and non-growing cells expressing a range of TetA levels (arrested). Interestingly, TetA expression in arrested cells occurred during a variable period of slow growth that preceded growth arrest (moribund). Although the coexistence of recovered and arrested cells has been explained by stable states corresponding to high and low TetA expression [10, 47], the high variability of TetA expression levels in arrested cells is not explained by models based on a bistable mechanism.

### Model

The dynamics of TetA and TetR are coupled to the metabolic state of the cell in two ways. First, in order to maintain stable protein concentrations, the rate at which all proteins are produced must scale with the rate (per unit mass) of cell growth *λ*, otherwise these proteins would be diluted [12, 25, 27, 34, 37]. This is true independently of resource allocation. Second, the cellular ribosomal content is adjusted in response to changes in nutrient or drug conditions, and this comes at the expense of the expression of non-ribosomal proteins, including TetA and TetR [19, 32, 33, 44] (Fig. 1D). According to the proteome partition framework by Scott et al., the fraction of the proteome composed of ribosomal proteins, denoted *ϕ*_*R*_, shows two linear relationships depending on whether growth is changed by varying drug or nutrient conditions. In the absence of drug, varying nutrient conditions results in ribosomal content increasing linearly with the cell growth rate *λ*:

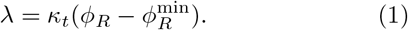

Here, *κ*_*t*_ is the effective rate at which ribosomes translate new proteins, and 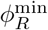 is the minimum ribosomal fraction needed to support growth. Meanwhile, under fixed nutrient conditions, varying the concentration of a translation-inhibiting drug results in ribosomal content decreasing linearly with the cell growth rate, to compensate for changes in translation efficiency:

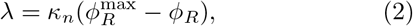

where *κ*_*n*_ is a parameter reflecting nutrient quality and 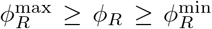. The remaining fraction *ϕ*_*Q*_ = 1 − *ϕ*_*P*_ − *ϕ*_*R*_ is called the *Q*-sector.

A variable fraction of the proteome, denoted *ϕ*_*P*_, which includes TetA and TetR, is modulated to counterbalance the changes in ribosomal content, following that *ϕ*_*P*_ = *λ/κ*_*n*_. Given the *P* -fraction of the proteome allocation in the absence of drug *ϕ*_*P*,0_, the ratio between the rates of expression of *P* -sector proteins in the presence and absence of drug obeys

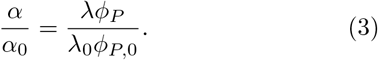

The factor *λ* captures the overall faster production/dilution rate, and since *ϕ*_*P*_ is also proportional to *λ*, the expression rate *α* scales with *λ*^2^. The dynamics of a protein *x* in the *P* -sector can then be written in the form (see Appendix A)

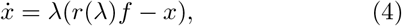

where *r*(*λ*) = *λ/λ*_0_ is a coarse-grained parameter representing the resources allocated to this gene and *f* = *α*_0_*/λ*_0_ is a drug-independent parameter which includes other processes modulating expression of this gene, such as regulation. The second term in Eq. (4) comes from protein dilution due to cell growth.

Our approach treats the coarse-grained variable *r* dynamically and assumes that *extrinsic* variation in *r* is the dominant source of noise. We view *r* as a hidden degree of freedom in the expression dynamics and ask the following question: Can a simple model for the *r* dynamics which is consistent with Eqns. (1) and (2) produce both the bimodal cell fates and the range of slow-growth phenotypes with variable expression observed in the experimental data? Other models that consider resource allocation to be in equilibrium fail to identify slow-growing states [47], but models considering the dynamics of resource allocation have shown promise by recapitulating other relevant metabolic interactions [19, 26].

We assume no transcriptional regulation during the course of the drug response, hence we set *f* = 1 for all genes. This assumption is valid while intracellular tetracycline concentration remains high, inactivating TetR and preventing repression of TetA production. Since recovered cells eventually reduce intracellular tetracycline concentrations enough for regulation to become relevant again, our model is not meant to predict the precise expression levels of recovered cells. Rather, the goal is to explain the emergence of phenotypic diversity and the trajectories of slow and non-growing cells where intracellular tetracycline remains high.

Let ***x*** = (*x, y, z*, …) be a vector of protein concentrations and ***r*** = (*r*_*x*_, *r*_*y*_, *r*_*z*_, …) the corresponding resources. For a given growth rate, we model (***x, r***) with linearized dynamics,

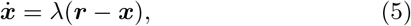

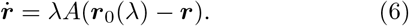

Here, ***r***_0_ is a growth rate dependent fixed point, which is chosen so that steady-state protein concentrations are consistent with Eqns. (2) and (3) for fixed *λ*. The matrix *A* specifies the coupling between the resource allocations for each gene, which do not necessarily evolve independently. This is a free parameter in our model, but as we explain below, our conclusions are not sensitive to the details of *A*.

The rationale for the factor of *λ* in the resources ***r*** equation is that ***r*** also represents concentrations of stable molecules, which are therefore subject to dilution just like protein concentrations ***x***. In Appendix C, we consider alternative interpretations of ***r***, illustrating that this assumption is not strictly necessary.

We focus on the case ***x*** = (*x, y, z*) with (*x, y*) representing resistance proteins (such as TetA and TetR) and *z* is a constitutively expressed gene in the *Q*-sector, which we include for comparison. To complete our model, we specify how the growth rate depends on the internal drug concentration and how the internal drug concentration depends on the expression of resistance proteins, which introduces a feedback from *x* to the growth rate. The growth rate *λ* depends on the effective translational capacity 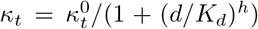, where *d* is the internal drug concentration, which is assumed to bind ribosome. Therefore, from Eq. (1)

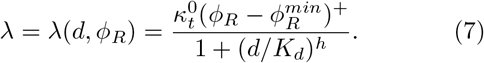

The drug-free growth rate 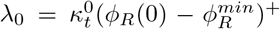 is determined by the resource allocation fractions and the drug-independent translational capacity 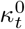 (see Appendix). The internal dynamics of the drug are

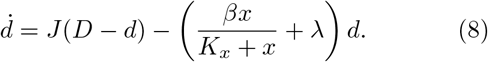

where *D* is the external drug concentration, *J* is the diffusion rate and *βx/*(*K*_*x*_ + *x*) is the rate at which the resistance protein removes the drug, given by Michaelis-Menten kinetics.

The time-scales of our model are *κ*_*n*_, 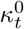, *β, J* and *λ*_0_. We set 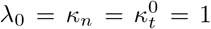, motivated by the fact that these are all the same order in growing cells. Therefore, time is measured in units 1*/λ*_0_. Henceforth, the concentration scales *K*_*d*_, *K*_*x*_, and *D*, as well as the time-scales *β* and *J* will all refer to the non-dimensionalized parameters. Since *x, y* are *P* -sector proteins and *z* is in the *Q*-sector, we have

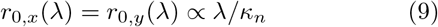

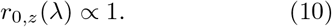

For simplicity, we set the constant of proportionality to be one, as this constant does not influence our key conclusions. These expressions are selected to ensure that as *t* → ∞ the scaling relations between growth and resource allocation fractions are respected.

### Expression dynamics following an abrupt drug exposure

We use the model to study how a growing cell responds to an abrupt change in the external drug from 0 to *D >* 0. Prior to exposure, transcriptional regulation determines the dynamics the initial conditions, where TetR is maintained at a level high enough to completely repress expression of TetA [42]. As mentioned above, our assumption is that regulation becomes irrelevant following exposure, since the drug saturates TetR [43]. This motivates the initial conditions *x*(0)*/K*_*x*_ = *r*_*x*_(0) = 0, *y*(0) ≈ *r*_*y*_(0).

We now turn to the coupling matrix *A*. We conjecture that the important feature of this matrix *A*, in addition to having positive eigenvalues, is that the coefficient of *r*_*y*_ in the *r*_*x*_ equation is negative. This reflects the fact that the two proteins share resources, since they are in the same sector of the proteome, so that an excess of *y*-resources, meaning *r*_*y*_ − *r*_0,*y*_ *>* 0, should result in a positive contribution to the production rate of *x*-resources.

For our simulations, we set

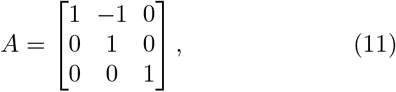

which contains only the interaction that is relevant the the production of *x*, which is the resistance protein that interacts directly with the drug. However, the relationship between *x* and *y* needs not be asymmetric in order to produce experimentally observed expression level correlations (see Appendix C). In using this form of *A*, we also assume that *z* is only coupled to the *P* -sector genes via its dependence on the fixed point (Eq. 10).

In our simulations with *A* as given above, we observe a drug-dependent critical value of 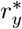 for the initial *y*-resources. Below this threshold cells become trapped in a slow manifold with a slow growth rate, while above this threshold cells quickly recover to their pre-drug growth rate (Fig. 2A). Trajectories over a range of *r*_*y*_(0) show a similar pattern of variation to the experimental data (2B). TetR (*y*) levels are spread out over a wide range in slow-growing cells, but they are instead concentrated around the fixed point in growing cells. We emphasize that the slow-growing cells are not at a fixed point, but will eventually recover on a slow time-scale.

**FIG. 2.**
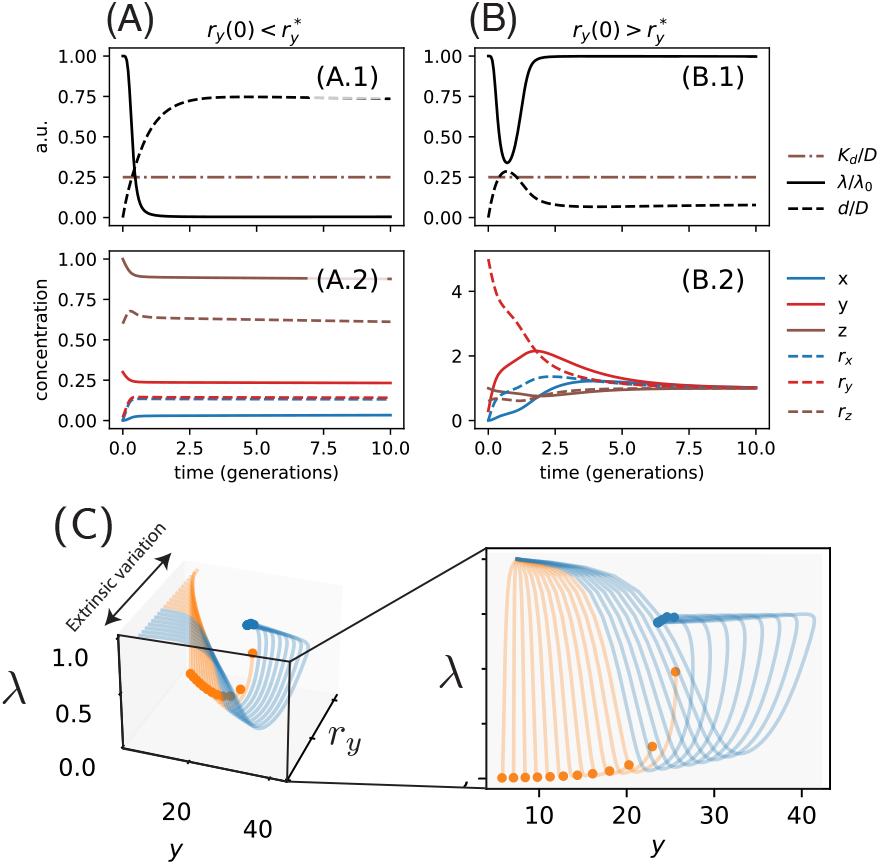
Examples of trajectories for values of *r*_*x*_(0) with initial *y*-resources *r*_*y*_(0) (A) below and (B) above the critical 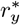 threshold. (A.1) and (B.1) show the growth rates and internal drug concentration. The brown line is threshold for drug to inhibit cell growth. (C) The dynamics for many different initial resource fractions *r*_*y*_ (0) in *x* − *r*_*y*_ − *λ* space and projected to the *x* vs. *λ* plane (right).

This model can also reproduce other characteristics of the experimental data, such as the correlation between TetA and TetR levels in slow-growing cells and the correlation between TetR and the constitutively expressed protein (Fig. 3). It is important to note that we have not explicitly modeled the initial variation in resources, which are prescribed by the dynamics prior to exposure. This requires a more careful consideration of the interplay between regulation and resources (Appendix B). When regulation is present, it is possible to have high levels of variation in *r*_*y*_ with virtually no variation in *y*, even though the production rate of *y* scales with resource concentration.

**FIG. 3.**
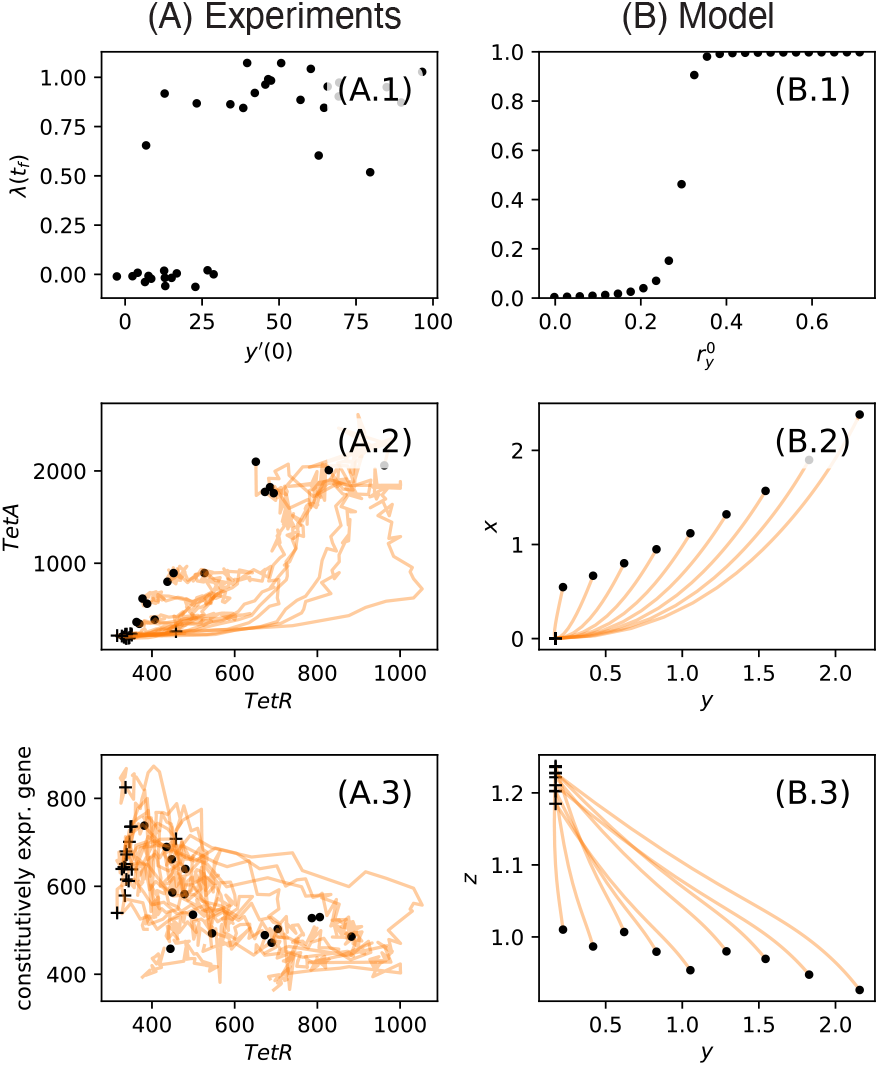
(A) Experimental data. (A.1) Final growth rate as a function of the initial TetR velocity. Trajectories of (A.2) TetA and (A.3) Constitutive protein (RnaI) vs TetR in arrested cells. The orange crosses mark the initial values of each lineage prior to exposure. (B) The same plot for simulations of our model.

### Critical resource for a threshold model

Here, we study the dependence of the critical resource 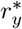 on the external drug concentration *D* (Fig. 2). The full system of ODEs for (***x***, *d*) is difficult to study analytically. However, in the spirit of work on gene regulatory networks and other nonlinear systems, we are able to gain some insight by treating nonlinearities as thresholds [2]. This approach does not give accurate expressions for the trajectories, but it does help understand the mechanism leading to the divergent cell fates and the scaling of the critical resource with external drug. This critical resource value can be viewed as an escape “velocity” for TetR, since the initial production rate is proportional to *r*_*y*_(0).

We first note that the fate of a cell is determined by two competing time-scales: First, there is the time *τ*_*d*_ for the drug to inhibit growth. Second, there is the time *τ*_*x*_ for TetA to reach high enough levels to overcome the influx of drug. A cell will recover when *τ*_*x*_ *< τ*_*d*_. We then suppose that the exponent *h* in the drug inhibition of growth is large enough, so that we can approximate growth *λ* as depending on a threshold in the internal drug concentration *d*. This means that for *t < τ*_*d*_ growth does not change significantly. Although in reality cell growth varies substantially with the internal drug concentration around the threshold, this approximation still allows us to capture the scaling behavior of the critical resource 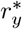 in *D*. Since *λ*(0) = *O*(1) and the external drug concentration is large *D* ≫ 1, an initial constant influx of drug results in the internal drug concentration increasing linearly in time *d*(*t*) ≈ *JDt*. Meanwhile, TetA concentration grows quadratically as *x*^′^(*t*) = *λ*^2^*r*_*y*_(0)*t*^2^*/*2 + *O*((*tλ*)^3^), where we have set *y*(0) = *r*_*x*_(0) = 0. Considering that TetA production does not significantly alter the influx of drug until it reaches high concentrations, the influx of drug balances the removal via TetA at a value *x*^*^, satisfying the equation

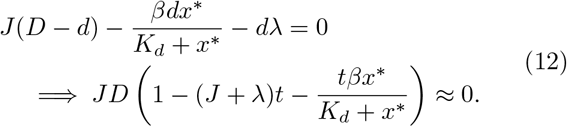

Thus, *x*^*^ is *O*(1) in *D*. Note that if *τ*_*d*_ *< τ*_*x*_, then *d*(*τ*_*d*_) ≈ *K*_*d*_, implying that *τ*_*d*_ ≈ *K*_*d*_*/JD*, while *x*(*τ*_*x*_) = *x*^*^. The condition for *τ*_*x*_ *< τ*_*d*_ is therefore

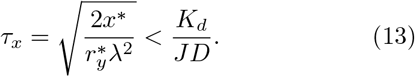

It follows that the critical value is 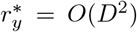. This scaling is consistent with our numerical simulations (Fig. 4A). The specific scaling with *D* is sensitive to the structure of *A* (unlike the correlations in Figure 3) and could therefore be helpful in determining the structure of the coupling matrix from data (see caveat below).

**FIG. 4.**
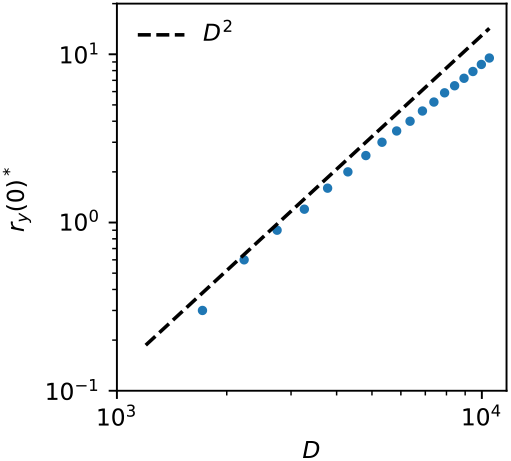
The scaling of the critical resource concentration 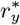 as a function of the external drug concentration *D*. All other parameters are the same as in Fig. 1.

## Discussion

In this study, we proposed a simple model that couples resource allocation dynamics with the expression of resistance genes and the dynamics of accumulation of intracellular drug. The main motivation for our model was to identify an alternative mechanism for the emergence of phenotypic heterogeneity during drug responses, expanding on previous models accounting for bistable cell fates [1, 4, 5, 10, 13, 15, 28, 40, 48]. In particular, our model provides an example of a system where slow trajectories generate transient bimodality that disappears in the long-time limit. While our model simplifies several aspects of cellular physiology – such as neglecting the regulation of the drug response – it provides a tractable framework for understanding how cellular heterogeneity arises from fundamental constraints on resource allocation.

The slow trajectories emerge from a diverging timescale, which results from the assumption that the linearized derivatives of all concentrations are proportional to the single-cell growth rate. As ribosomes become saturated, the growth rate vanishes and the relaxation timescale diverges. We expect this is a general feature of physiological systems under stress, where cell responses are induced under inhibition of cell growth, and so our model should be of wide interest beyond the application to the *tet* system.

The nature of the resources *r* allocated for gene expression can be interpreted as any quantities that change dynamically and modulate the expression rate. In the case of our experiments, where TetR and TetA share resources, but TetA initial resources are low while TetR are high. The resources in question can be interpreted as the availability of mRNA or other proteins involved in transcription or translation prior to exposure. TetA and TetR resources are shared because they belong to the same fraction *P* of the proteome, but TetA mRNA levels are kept low prior to drug exposure because of regulation by TetR. The resources available to TetR then represent a measure of the resources available to the whole sector of the proteome.

Interestingly, using order-of-magnitude estimates of the parameters, our model successfully reproduced key experimental observations, including distinct cell fates and the existence of a critical threshold in TetR production velocity that determines whether the pre-drug growth rate is restored on a fast timescale [42]. In principle, by examining how this critical velocity scales with the external drug concentration, one can infer properties of the underlying coupling matrix. In practice, however, this is difficult due to other sources of variation that make it challenging to define a clear critical velocity from the experimental data. Future work could extend this model by explicitly incorporating stochastic fluctuations in gene expression and interactions with other cellular pathways involved in the antibiotic response.

## Acknowledgments

Levien and Schultz acknowledge support from NSF DMS-2527337. Schultz acknowledges support from NSF PHY-2412766. Levien acknowledges helpful discussions with Jie Lin and David Lacoste, and thanks the organizers of the 2025 Gordon Research Conference on Stochastic Physics in Biology for providing an opportunity to share preliminary results from this paper.

## Code and data availability

All the data and code used in this paper is available at https://github.com/elevien/AntibioticsCellFate.

## Appendix A: Model derivation

Let *x* denote the concentrations of a generic protein (e.g., TetR, TetA, constitutive protein). The time derivative of *x* is given by a production and dilution term [34]:

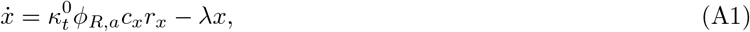

where *c*_*x*_ is a condition-independent, gene-specific constant, *r*_*x*_ is a dimensionless number, the fraction of resources allocated for translation for protein *x* (including mRNA and amino acids), 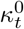 is the translational efficiency in the absence of drug, and *ϕ*_*R,a*_ is the fraction of actively translating ribosomes. We set *c*_*x*_ = 1 since this can be absorbed into *r*_*x*_. *ϕ*_*R,a*_ is not to be confused with the ribosomal resource allocation fraction. Note that while the translational efficiency defined in Eq. (1) is drug dependent, 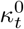 is a drug-independent parameter. Our model assumes that ribosomes are the growth limiting molecule and Eq. 8 follows from the formula 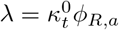.

There are different ways of incorporating noise into this model. First, we can consider noise in gene expression resulting from stochasticity in mRNA and protein production [34] – we will refer to this as intrinsic noise, although this includes the binding and unbinding of transcription factors to the TF bindings sites, which by some definitions is extrinsic. Since the coarse-grained variable *r*_*x*_ incorporates mRNA copy numbers, intrinsic noise will lead to fluctuations in *r*_*x*_ around a fixed point which is condition-dependent. We do not model this source of noise, but rather focus on extrinsic variation in the fixed point itself.

Instead of trying to come up with a mechanistic model of ***r***, we take the approach of imposing the constraint that the proteome partitions are respected in fixed conditions. To this end, let

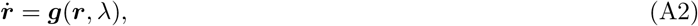

where *g* is some nonlinear function depending on the growth rate (equivalently, the fraction of actively translating ribosomes; that is, ribosomes not bound to the drug). We assume that *g* has a fixed point

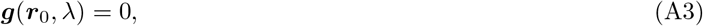

where ***r***_0_ respects Equations. (1) and (2), meaning for *P* -sector proteins *r*_0_ → 0 as the drug causes *λ* → 0. We linearize around this fixed point to obtain

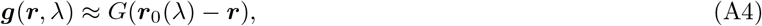

where *G* = (*D****g***)(***r***_0_, *λ*) is the Jacobian of ***g*** at the *λ* dependent fixed point ***r***_0_(*λ*)

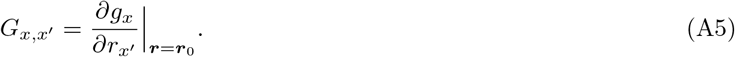

We are using *g*_*x*_ to represent the *r*_*x*_-component of ***g***.

In the main text, in order to reduce the space of possible models, we make the assumption that *G* depends on *λ* only by scaling: There is a *λ*-independent matrix *A* such that *G* = *λA*. This is motivated by the fact that the dilution time-scale is set by *λ*. If “resources” are proteins, such as ribosomes or RNA-polymerases with slow degradation, it is natural to model their dynamics in this way. An alternative model would be to take *G* = (*λ/λ*_0_)^*ω*^*μ*, where *μ* is a *λ* independent rate and *ω* is some power-law constant.

## Appendix B: On the initial variation in *y* and *r*_*y*_

Here we address the question of how to interpret the initial conditions biologically. We first discuss the high variability in *r*_*y*_. In experimental data and in our simulations the CV of *r*_*y*_(0) is on the order of 1, but the CV of *y*(0) is effectively zero. These may seem like contradictory initial conditions because the production rate of *y* prior to exposure is scaled by *r*_*y*_(0). A potential explanation is nonlinearity in the production rate. Let *λr*_*y*_*f* (*y*) be the production rate where *f* (*y*) is a nonlinear function that captures the influence of regulation. The natural model for *f* (*y*) is

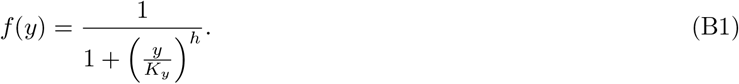

If the nonlinearity is strong enough then large variation in *p*_*y*_ will have little effect on the solution to

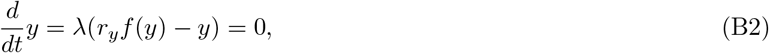

which will be *y* ≈ *K*_*y*_.

Whether such a strong nonlinearity is present depends on the details of the expression dynamics and we leave it to future work to explore whether this is reasonable for the *tet* system. It is possible that the interplay between the resources and regulation is more complex and requires considering feedback between the expression levels and the resource dynamics. Finally, note that whether we put the nonlinearity in the equation for *y* vs. the equation of *r*_*y*_ is a question of how we interpret *r*_*y*_ and will be pursued in future work.

## Appendix C: Sensitivity to model details

Here, we explored the sensitivity of our results to the details of the model, focusing on the matrix *A* and the dependence of the resource dynamics on the growth rate.

### 1. Thecoupling matrix *A*

We study how sensitive our results are to the choice of *A* by considering a parametric family of matrices,

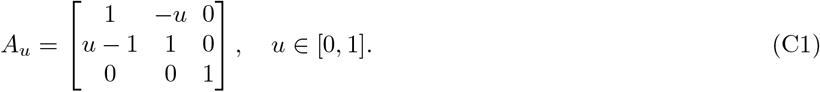

By modulating *u* between 0 and 1 we interpolate between the case where *y* follows (*u* = 0) and *x* follows *y* (*u* = 1). Hence, the matrix used in the main text is *A* = *A*_1_. For each value of *u*, we perform simulations over the same range of initial conditions used in the main text and obtain the following summary statistics:

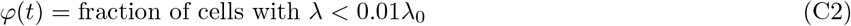

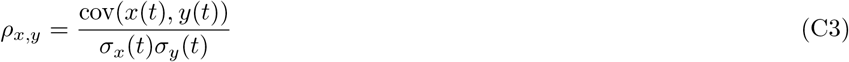

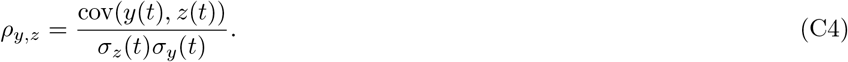

These are respectively the fraction of arrested (or slow-growing) cells, the correlation coefficients of *x* and *y* among arrested cells, and the correlation coefficient of *y* and *z* among arrested cells.

In Fig. S1 (A) we show *ρ*_*x,y*_ and *ρ*_*y,z*_ as a function of *u*. Above *u* ≈ 0.2 the summary statistics appear to be largely independent of *u* and consistent with experimentally observed values. We have also shown the values of these statistics for the data. In Fig. S1 (B) we have shown the *φ* as a function of time for different *u* values.

### 2. Growth rate dependence of resource dynamics

We next sought to understand how sensitive our results are the dependence of the resource dynamics on the growth rate. To this end, we simulated a generalization of the model given by

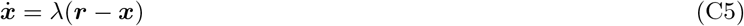

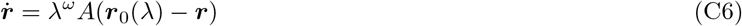

where the new parameter *ω* allows us to interpolate between a regime where the dynamics of the resources is not dependent on the growth rate (*ω* = 0) and the model simulated in the main text. We performed the same analysis as was done for *u* in the previous subsection, and the results are shown in Fig. S1 (C) and (D). Interestingly, it can be seen that for small enough *ω* the arrested cells appear to never recover. Therefore, the finite time summary statistics are not sensitive to the specific scaling of the resource dynamics with growth rate, it seems to strongly effect the long-time dynamics and determines arrested cells are a transient state or now.

**FIG. S1.**
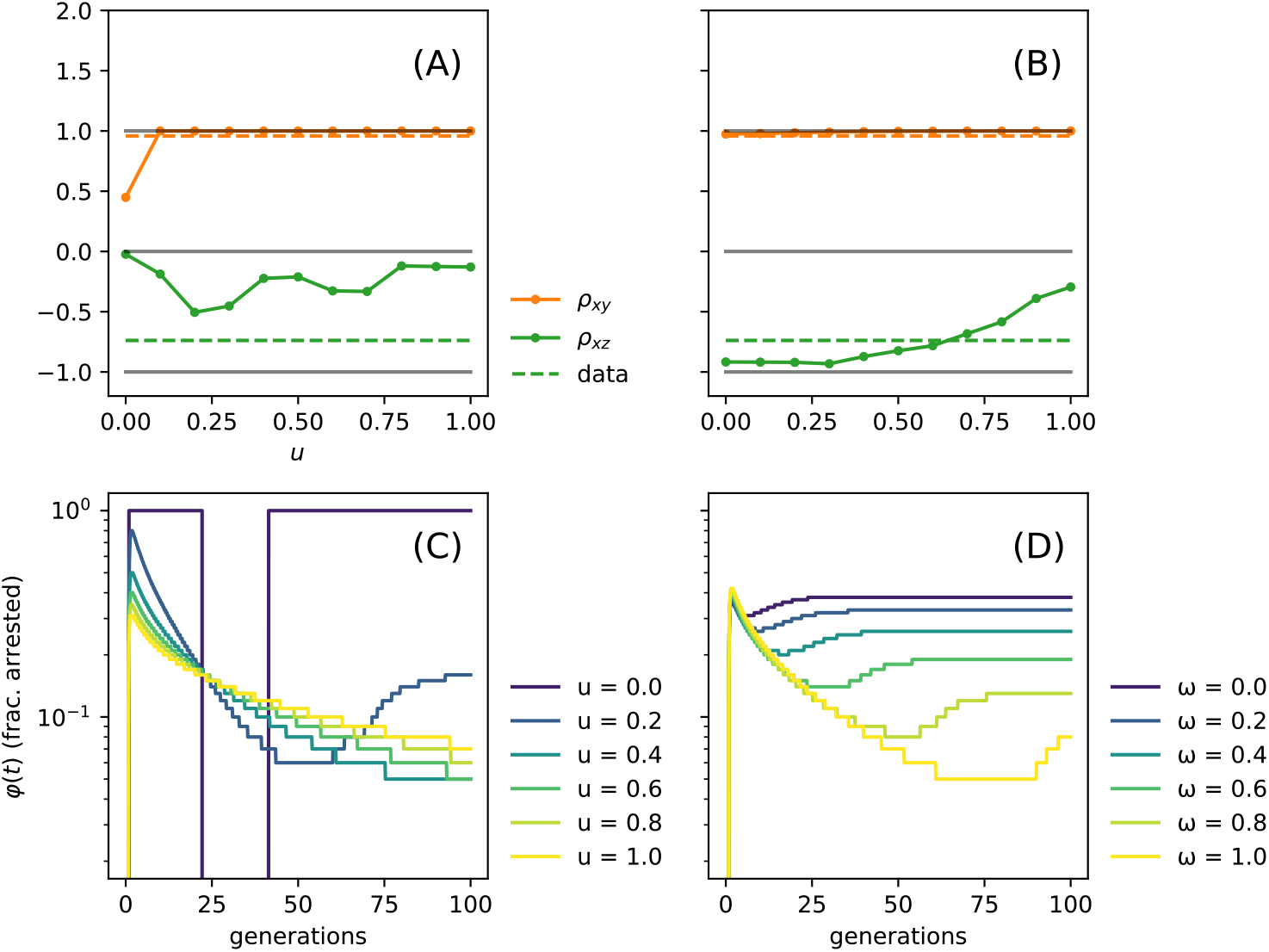
Sensitivity analysis of the model to variations in the resource coupling matrix parameter *u* and the growth-dependence parameter *ω*. (A) Fraction of arrested cells *φ*, and correlation coefficients *ρ*_*x,y*_ and *ρ*_*y,z*_, at a fixed time as a function of *u*. (B) Time evolution of *φ* for several *u* values. (C) Summary statistics as in (A) for varying *ω*. (D) Time evolution of *φ* for several *ω* values. All other parameters are identical to those used in the main text.

## Appendix D: Parameter values

We determined the non-dimensional parameter values given in Table I. To non-dimensionalized, divided *D, β* and *J* all the rate by *λ*_0_ = 0.015 – the growth rate in drug free conditions. The concentration scale was determined by the drug-free gene expression rate, which is ≈ 0.0003. Dividing this by *λ*_0_ yields a concentration scale *c*, which was used to obtain nondimensional *K*_*d*_, *K*_*y*_, *J* and *D*.

**TABLE I.**
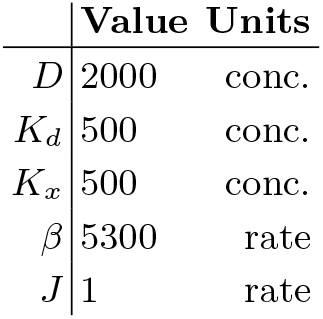
Nondimensional parameter values obtained with order of magnitude estimates from [47].

